# FLUID-CELL: Flow-enabled Light and Ultrastructural Imaging Device for Correlative Electron and Light Localization

**DOI:** 10.1101/2025.06.17.660254

**Authors:** Nicholas M. Rienstra, Steve Garvis, Juan C. Sanchez, Bryan Sibert, Elizabeth R. Wright

## Abstract

We present a novel microfluidic flow cell, or Flow-enabled Light and Ultrastructural Imaging Device for Correlative Electron and Light Localization (FLUID-CELL), that connects fluorescence light microscopy (FLM) and cryo-electron microscopy (cryo-EM) for advanced biological imaging and ultrastructural and structural analyses. The design of the FLUID-CELL features a precisely engineered microchannel that maintains native cell culturing conditions, supporting correlation and enabling real-time observation by FLM, as well as subsequent cryo-EM analysis. In this study, this device enabled practical FLM imaging over extended experimental periods, consistent sample handling, and the ability to perform correlative imaging. This capability connects dynamic cellular events imaged by fluorescence light microscopy with high-resolution ultrastructural data collected with cryo-EM. Our dual-modality approach streamlines the workflow and opens new possibilities for investigating the relationship between cellular function and molecular architecture at the nanoscale.

## Introduction

Cryo-correlative light and electron microscopy (cryo-CLEM) bridges the gap between fluorescence microscopy and cryo-electron microscopy (cryo-EM) imaging^1^. Cryo-electron tomography (cryo-ET) permits the creation of three-dimensional (3D) reconstructions of intact biological specimens in their native state^2^. Over the last several years, rapid advances in imaging technologies have enabled the visualization of biological structures at nanometer and sub-nanometer resolutions, making cryo-EM and cryo-ET crucial tools for investigating macromolecular complexes in their native states. Correlative light and electron microscopy (CLEM), combined with the sample thinning capabilities of cryo-focused ion beam (FIB) milling, provides a means for identifying biological targets essential for the directed milling of thick cellular samples to reduced thicknesses of ∼100-300 nm, which is required for cryo-EM and cryo-ET investigations. However, challenges in sample preparation persist, including maintaining biological integrity, preventing both biological and ice contamination, and controlling ice thickness^3,4^. These negative factors can impact prospects for achieving sample reproducibility and high-throughput data collection.

Several improvements for culturing prokaryotic and eukaryotic cells for CLEM, cryo-CLEM, and cryo-EM investigations have been introduced recently. EM substrate micropatterning is used to functionalize specific regions of a material, supporting cell adhesion and growth in defined patterns and shapes that conform to cell sizes and preferential growth characteristics^5–11^. The use of micropatterning also facilitates complex cryo-CLEM workflows, where maintaining cells in targeted locations on a substrate can maximize the number of relevant biological events and increase throughput. In parallel, 3D printing technology has advanced, and the range of available printing materials now includes biologically compatible materials^12^. These advancements have enabled rapid prototyping capabilities, allowing for the swift design, printing, testing, and revision of multiple iterations of a tool component in a single day within a single lab. Several cryo-EM-compatible 3D-printed tools and cell culture platforms have been introduced to facilitate small-volume workflows, reduce sample and substrate manipulation, and support multi-cellular communities^13–15^. These developments have reduced some of the challenges associated with multifaceted cell culture and cryo-CLEM studies, thereby making them more accessible to new investigators.

Microfluidics has also seen remarkable advancements over the past two decades. Microfluidic-based equipment and workflows enable precise, streamlined, and small-volume sample preparation and manipulation. These systems have emerged as solutions for automating and standardizing cryo-EM sample preparation for single-particle cryo-EM^16–21^. A key advantage of microfluidics is its ability to integrate and control multiple experimental variables. These variables include simple time-resolved reagent mixing, gradients of chemical stimuli, and hydrodynamic forces, thereby supporting efforts to examine interactions between mechanical forces and biochemical signals, such as those involved in nano-scale molecule-to-molecule interactions or larger-scale factors that regulate cell growth and biofilm development. Early pioneers in light microscopy have demonstrated the potential of microfabricated devices that control fluid dynamics for various biological applications, including culturing bacteria, neurons, and other cells, thereby supporting live-cell imaging to monitor the spatial and temporal dynamics of communities, cells, organelles, and macromolecules^22–25^. Thus, the incorporation of 3D-printed microfluidic devices for live-cell imaging offers the potential to further extend the capabilities of CLEM imaging.

CLEM workflows are used to locate and image specific biomolecules, and cellular structures present within target cells. Merging microfluidics with CLEM will enhance imaging efficiency, bridging the gap between dynamic, live-cell, physiological observations made with light microscopes and high-resolution, static cryo-FLM and cryo-EM imaging. Many technological advancements have streamlined cryo-CLEM workflows, including automated cryo-stages and integrated fluorescence modules, which have reduced the need for manual intervention when correlating imaging data^1^. The ability to perform simultaneous or sequential imaging helps minimize artifacts, ensuring that fluorescent signals align accurately with the ultrastructural details captured by electron microscopy (EM). These integrative approaches have been crucial in uncovering complex molecular interactions that were unseen due to limitations in imaging techniques.

The convergence of microfluidics, correlative light and cryo-electron microscopy (cryo-CLEM), and cryo-electron tomography (cryo-ET) represents transformative methodologies for high-resolution structural cell biology. Here, we present a microfluidic device, Flow-enabled Light and Ultrastructural Imaging Device for Correlative Electron and Light Localization (FLUID-CELL), that facilitates direct live-cell imaging of bacterial cultures using fluorescence light microscopy (FLM). We demonstrate that at appropriate time points during an experiment, the fluid flow to the system can be stopped, the EM grids removed, and the EM grids negatively stained or vitrified for conventional EM or cryo-EM analysis, respectively. Using correlation microscopy software, such as CorRelator^26^, specific fluorescent or electron-dense fiducials, biomolecules, and cellular structures can be sequentially targeted and imaged for correlative, contextual, and high-resolution imaging studies. In summary, the FLUID-CELL device facilitates the physiological preparation and live-cell imaging of multicellular communities for cryo-CLEM investigations.

## Results and Discussion

The FLUID-CELL unit was designed and created to encapsulate an EM grid for fluorescence light microscopy (FLM) imaging under physiological conditions, in a sterile manner, and to facilitate its retrieval for analysis by negative stain electron microscopy (NSEM) or cryo-electron microscopy (cryo-EM).

## FLUID-CELL chamber design and fabrication

The flow chamber was fabricated using a Formlabs Form 3+ SLA 3D printer. The CAD drawings of the individual FLUID-CELL components were uploaded to the Formlabs Preform software. The software then guided the production of the flow cell components from stereolithographic resins (Figs. 1 and 2). The two main upper and lower body parts were printed using Clear V4 resin at a layer height of 25 µm, which lock together to form a sealed, leak-proof chamber. Luer-Lok port adaptors were integrated into the design to support the rapid exchange of tubing and fluids, with tolerances that ensure uninterrupted fluid flow.

**Figure 1.**
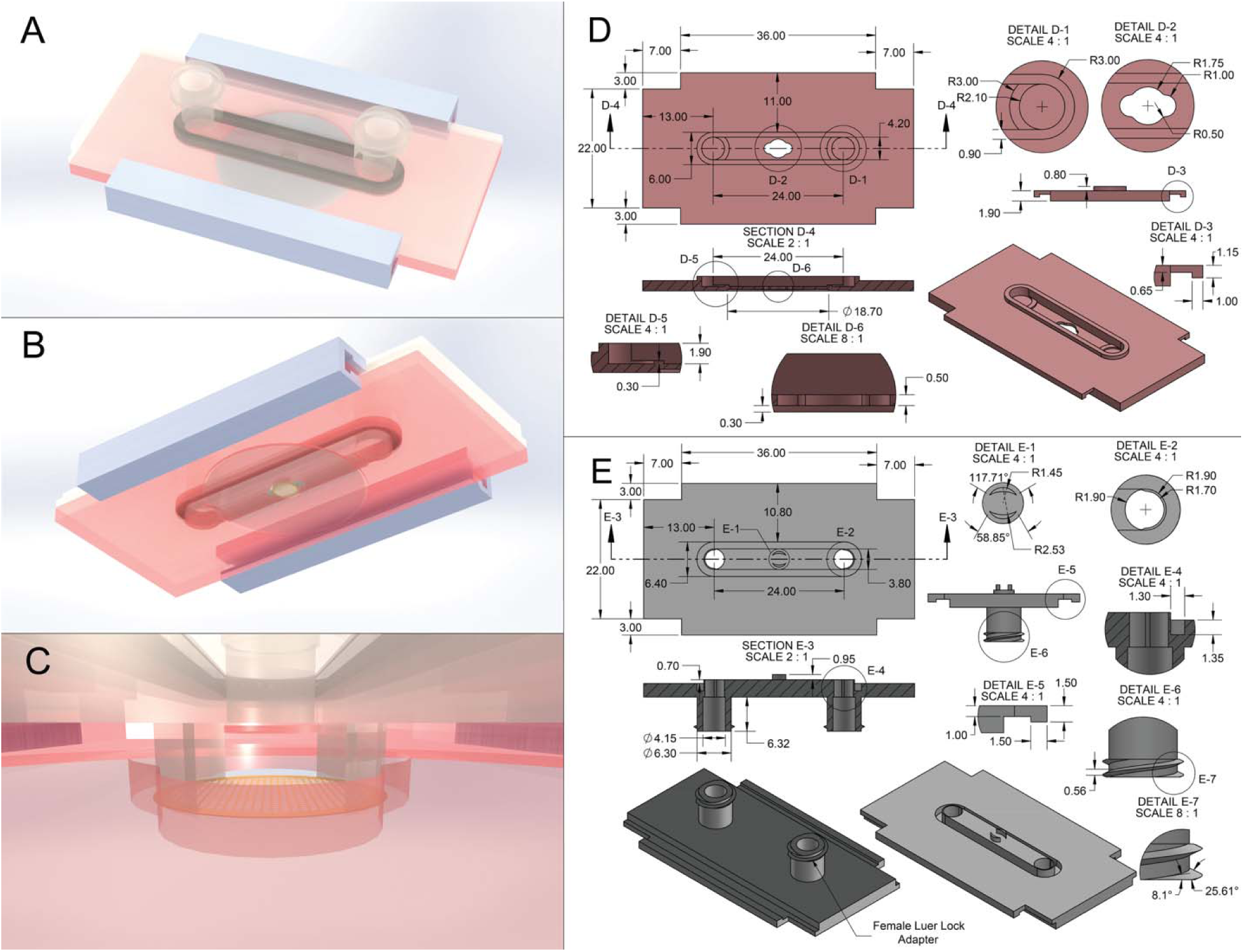
Design of the primary FLUID-CELL components. A) An overhead view of the system, with the gaskets (in black) and the Luer-Lok adaptor inlet and outlet ports (in transparent gray) visible. B) A view underneath the flow cell, the coverslip, and a centered grid is visible. C) A view of the flow cell from within the chamber. (D) Technical drawing describing the bottom section of the flow cell. Details of all relevant dimensions. (D-1) The base of the flow cell well shows the thickness of the chamber wall that compresses the gasket. (D-2) Receiving well for an EM grid. The central circle was chosen to allow for a small tolerance in grid placement. The smaller spaces to the left and right of the central circle are provided to allow for the development of flow progression before encountering the grid. (D-3) Receiving well for external clamping. (D-4) Cross section of the flow cell, along the path of the flow chamber. (D-5) Close up to the inflow port. Shows the height of walls surrounding the flow chamber. (D-6) Details the grid well depth as well as the depth of the coverslip well. (E) Technical drawing describing the top section of the flow cell. Details of all relevant dimensions. (E-1) Extruded ‘Bananas’ which keep the grid in the focal plane of the microscope, on a glass coverslip. Geometry was determined from the intersection of two arcs, one from the center of the grid well with a radius of 1.45 mm. The second was chosen as the center point of the previous edge, with a chosen radius of 2.53 mm. (E-2) Note the connections from the bottom of the inlet ports to the interior wall of the ports and the main body. (E-3) Cross section of the flow cell, showing the dimensions of the ports and the extrusion of the ‘bananas.’ (E-4) Shows the size of the gasket reception well. (E-5) Details the receiving well for clamping. (E-6) Shows the width of the threaded adaptor. (E-7) Shows rise and turn of the thread.

**Figure 2.**
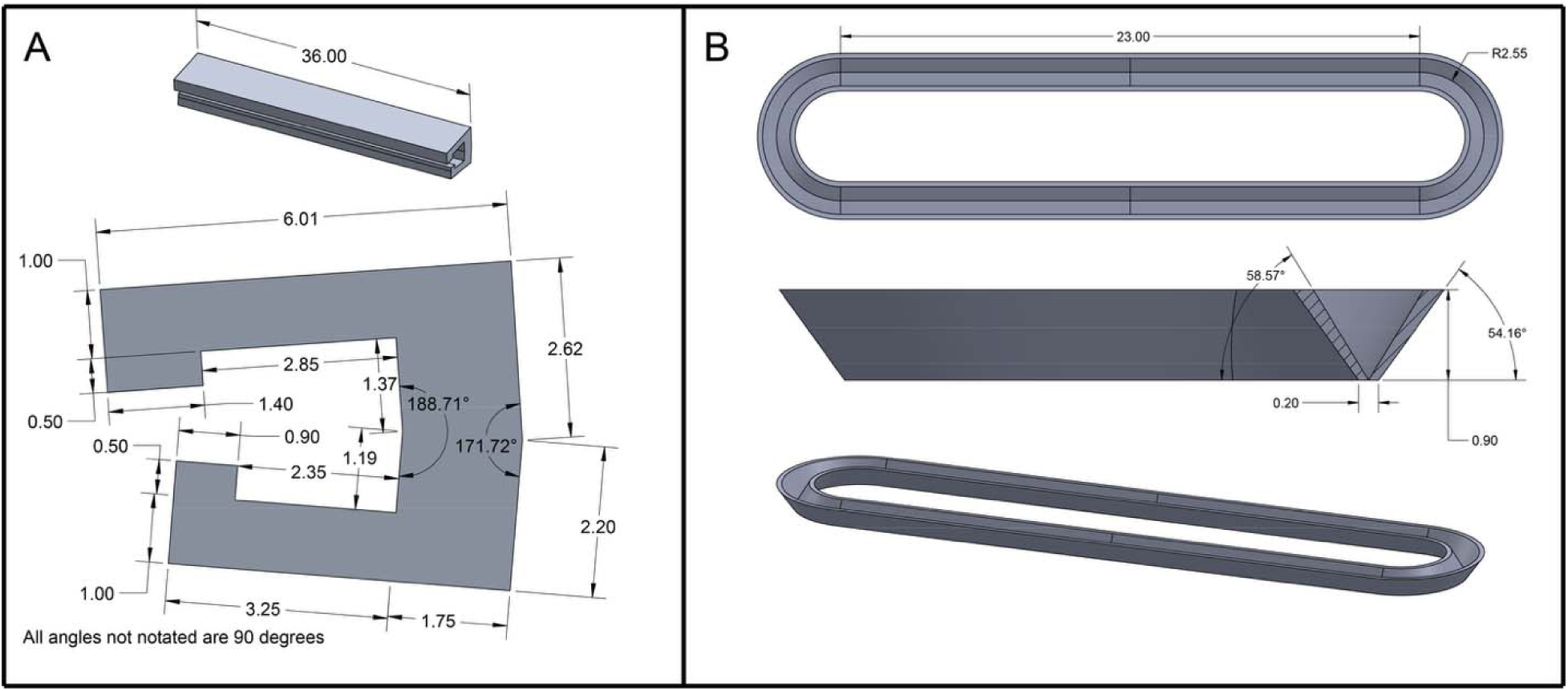
Design of peripheral FLUID-CELL components. (A) Technical drawing of the external securing bars and their dimensions. The bars are made of Durable V2 resin and are used to provide external pressure on the flow cell, compressing the gasket and sealing the flow cell chamber to make it leakproof. (B) Technical drawing of the sealing gasket and its dimensions. The sealing gasket is made of Silicone 40A resin and serves as a compressible seal between the two flow cell chamber halves, making it leakproof.

External securing bars were designed and dimensioned to fit the grooves of the FLUID-CELL (Fig. 2A); however, to produce the required pressure on the FLUID-CELL for compressing the gasket and sealing the chamber, dimensions and design were adjusted through prototyping. The revised dimensions are noted in Figure 2A. The bars are made of Durable V2, a more flexible and stronger material than the material of the FLUID-CELL, at a layer height of 50 µm. Deformations of the bars were drastically larger than those of the FLUID-CELL at the contact surface. This led to a pre-flexure of the bars, causing increased pressure that resulted in the expected deformations of the bars. Viscoelastic silicone 40A, at a 100 µm layer height, was used to manufacture gaskets that sealed the FLUID-CELL chamber (Fig. 2B), which encircled the region containing the EM grid and the flow inlet and outlet (e.g., Luer-Lok ports). Like the external securing bars, the gasket design and manufacture were an iterative process that began with a geometrically perfect design. Through iteration, it was found that a tapered cross section allowed for optimal dimensioning and placement in the FLUID-CELL chamber. Additionally, the overall sizing was selected to ensure a wall thickness that would be structurally sound during post-processing and usage of the gasket, while allowing for the smallest possible gasket for ease of compression.

We further employed concepts of additive manufacturing to extend the design and function of the FLUID-CELL. An example of a design extension is the ‘bananas’ that gently hold the grid in place on the glass coverslip (Figs. 1C, 4A, and 5A). These small, unique features would present significant challenges to manufacturing using traditional fabrication methods, which often require more standardized shapes and structures. In contrast, additive manufacturing, specifically 3D printing, enables the creation of these intricate components with precision and ease. By employing 3D printing technology, we are not only able to design the ‘bananas’ to hold the EM grid securely but also to optimize their shape and size. This minimized the surface area needed to be kept clear of the flow path, thus enhancing the overall efficiency and performance of the final FLUID-CELL. This level of design freedom underscores the advantages of additive manufacturing, enabling us to push the boundaries of what is possible with this device and future 3D-printed products.

## FLUID-CELL chamber operation, testing, and nanobead flow analyses

We used a programmable Gilson MINIPULS 3 peristaltic pump to manage media flow through the device. Flow rates during experiments were maintained at 200 µL/min but could be adjusted as needed. The MINIPULS 3 system enabled precise control of volumetric flow rates by changing the revolutions per minute (RPM) of the roller head. Our system used a 10-roller MP pump head to minimize pulsations in the flow. With the MP pump head, we used a linker portion of peristaltic tubing with two retaining stops (Gilson: F117942). The stops ensured that the tubing was correctly tensioned when fitted into the head. Compatible tubing, purchased from the Gilson company, was inserted into the peristaltic pump and connected to the flow system between the inlet flask and the device (Fig. 3A). Using tabulated information from Gilson correlated to the head angular velocity and tubing size, volumetric flow rates were calculated to compare with those generated by the pump. During fluorescence light microscopy imaging using the Leica DMi8, fluid flow was paused due to longer exposure times, which ensured that the bacteria remained more stationary in the optical path of the microscope.

**Figure 3.**
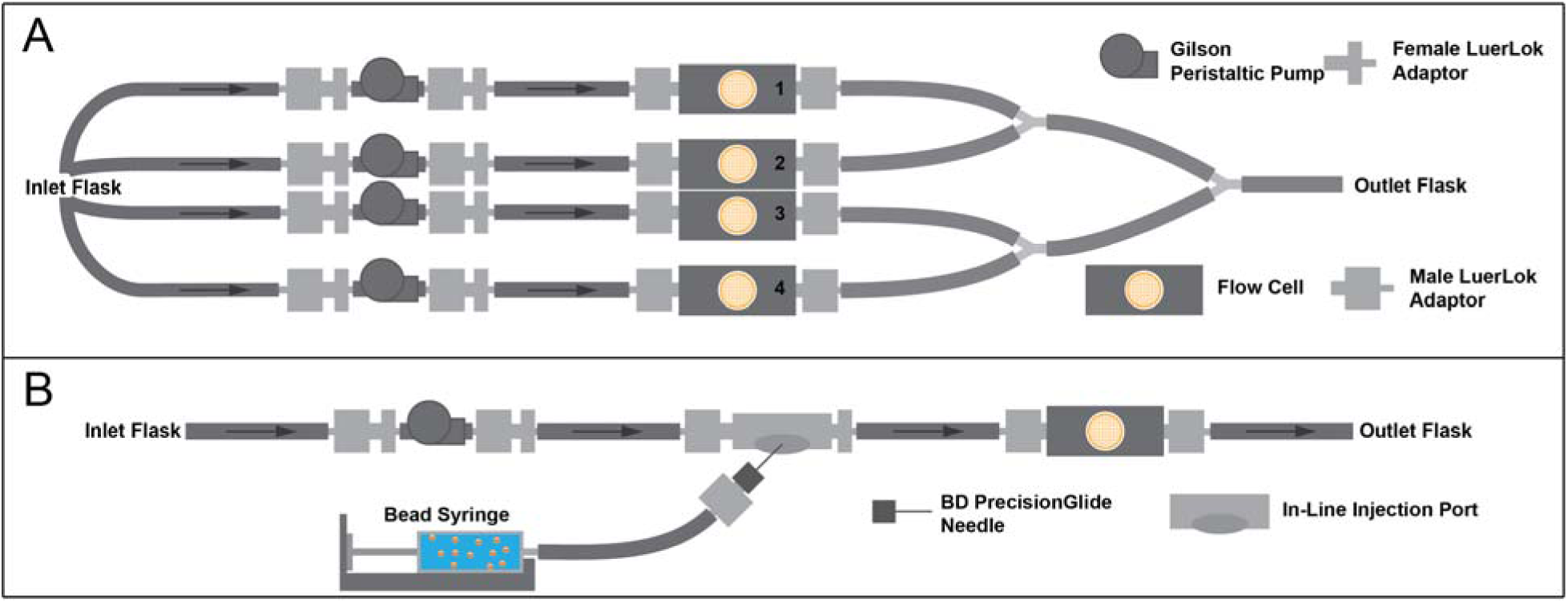
Diagram depicting flow cell setup. Diagrams are of the experimental configurations for bacterial cell culture (A) and nanobead flow experiments (B). (A) Four replicate FLUID-CELL devices were used in each experiment. However, other numbers of FLUID-CELL devices can be connected and used in similar configurations, depending on existing instrumentation. Once EM grids were inoculated, inlet flasks were filled with a rich PYE media, allowing the EM grids to be flushed with a fresh, sterile, and steady flow of media while microcolonies and biofilms developed. (B) The syringe with the nanobead solution was connected to the Gilson MINIPULS 3 peristaltic pump line far enough away from the flow cell and grid to allow for complete mixing between the two streams before reaching the observation point in the flow cell chamber.

The fluid flow characteristics were determined by flowing through the chamber 1.0 μm FluoSpheres Polystyrene Microspheres suspended in sterile water. The microsphere suspension was placed into a syringe coupled to a ChemYX Fusion 100X syringe pump (Fig. 3B). The Fusion pump’s line was connected to the MINIPULS 3 system line via an injection port. The connections were located a sufficient distance from the entrance to the flow cell to ensure proper mixing of the solution and consistency of fluid flow. The volumetric flow rates of the MINIPULS 3 and Fusion pumps were selected to simulate flow conditions during live experiments and control dilution of the beads. The mixed bead suspension was introduced into the microfluidic chamber, and time-lapse fluorescence light microscopy was performed using an inverted Leica DMi8 microscope equipped with the appropriate filter sets. Images and movies were recorded using Leica LASX software. The movies were analyzed using TrackMate software, which tracked the position of beads within the frame of the flow chamber (Fig. 4). Fluid flow tracking of the nanobeads in the collected movies provided velocity profiles and streamlines of the nanobeads across the EM grid set to various heights above the EM grid (20–400 µm) to simulate height variation that would exist in multicellular communities (e.g., bacterial biofilms). The beads travelled along predicted streamlines, creating a homogenous velocity profile across the center of the EM grid. These streamlines demonstrated a lack of turbulent or recirculating flow regions over the EM grid, indicating that the no-slip assumption made when calculating theoretical flow conditions was accurate, and that shear stress conditions at the grid surface were controllable through the volumetric rate of fluid moving through the chamber.

**Figure 4.**
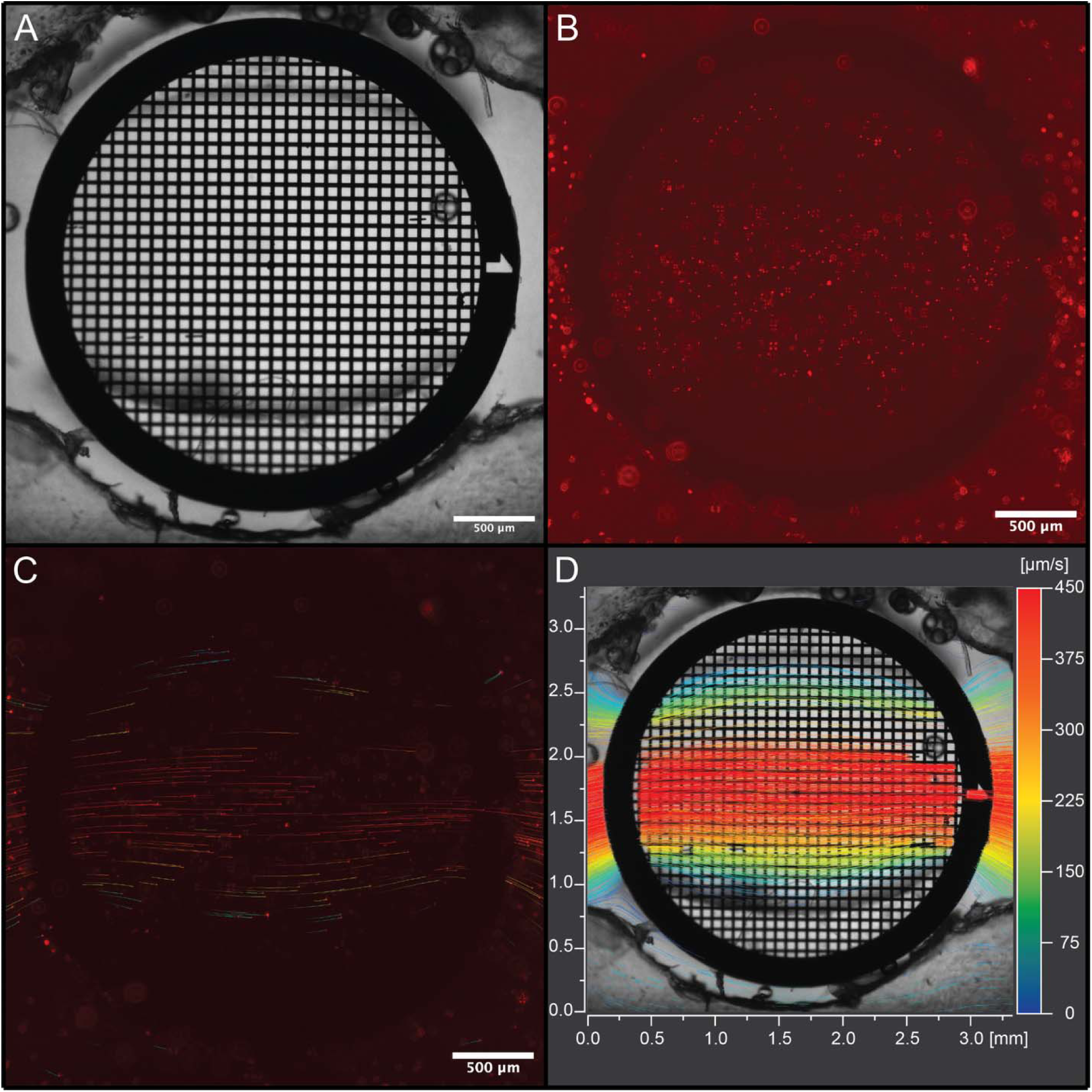
Velocity profile of the FLUID-CELL chamber with an EM grid in place. Velocity was calculated by passing fluorescent beads into the flow stream set at 100 µl/min and capturing a time-lapse image. Analysis was performed with FIJI software using the TrackMate plugin. (A) Field of view (FOV) data were collected using bright-field light microscopy. (B) View of this same FOV under Texas Red (TXR) fluorescence, with fluorescent beads visible moving across the top of the grid. (C) Single frame from a collected movie showing the flow paths that were processed from the movies, paths are colored based on a velocity gradient seen in D. (D) Collection of all flow paths from an entire data movie, depicts velocity gradients across the EM Grid as well as the observable streamlines that the fluid moves along. Scale bars of 500 μm (A-C).

To simulate shear conditions at the central region of the device surrounding the grid, the pump’s volumetric flow rate was adjusted. Due to the limitation of Reynolds’ numbers in microfluidic systems^27^, turbulent interior flow was not observed (Fig. 4). However, like aerodynamic experiments in wind tunnels, variations in flow can be achieved and observed through the manipulation of the pressure, density, and viscosity of the fluid media within the flow cell. Data analysis of the fluorescent nanobead experiments generated per-bead velocity profiles. The beads traveled consistently along predicted streamlines, which indicated that the additive geometry of the flow cell interior did not disrupt the smooth boundary condition assumption made during design (Fig. 4). These results are crucial, as uniform flow is necessary for studying biofilm formation under controlled shear stress conditions. The simulation of realistic flow can be used to study various biological specimens under different conditions, ranging from quasi-static fluids to variable dynamic flows and even non-Newtonian fluid flow (e.g., resembling blood flow). This versatility demonstrates that the flow cell can be tailored for various biomedical, industrial, and biological experiments.

## FLUID-CELL chamber operation with *Caulobacter crescentus* bacteria

To determine the applicability of the system for bacterial agents, we used *Caulobacter crescentus* strains CB15 and DH1210 (pleC::venus, divJ::mkate)^28^. The cells were cultured in standard peptone yeast extract (PYE) medium at 30 °C with shaking at 200 rpm. The bacteria were cultivated overnight and then diluted to an optical density (OD_600_) of 0.3, ensuring a healthy and active cell population. To simulate initial bacterial attachment, 200 µL of the diluted cell suspension was inoculated into the flow chamber and incubated for 16 hours at 30 °C. Following the 16 hr incubation, fresh PYE was pumped through the system. Fluorescence light microscopy images were taken at 0 hr of fluid flow to establish the initial colonization state (Fig. 5). Once flow was established within the system, most of the bacteria that had been inoculated into the flow cell chamber flowed freely to the waste flask, leaving behind individual or pairs of bacteria that had attached and remained on the EM grid surface. As the fluid flow continued, bacterial growth was monitored using time-lapse fluorescence light microscopy to track the development and maturation of the biofilm (Figs. 5 and 6). *C. crescentus* has a doubling time of approximately 90 minutes; therefore, images were taken at ∼60-minute intervals to capture a full range of cell growth dynamics. Parameters such as the rate of colonization, biofilm thickness, and spatial heterogeneity of the biofilm were monitored. These observations were then correlated with the previously obtained flow characteristics using a standard Ibidi flow cell (microslide VI.01) to confirm that this flow system functioned similarly and that bacteria could colonize the EM grid, developing a biofilm comparable to that in standard commercial flow cells (Fig. 5). It should be noted that the imaging data were not at synchronized time intervals between the Ibidi flow cell and the FLUID-CELL; this is due to the easier attachment of *C. crescentus* to the flat glass coverslip of the Ibidi cell. This medium is easier for the bacteria to attach to, leading to faster biofilm development than to the EM grid in the FLUID-CELL.

**Figure 5.**
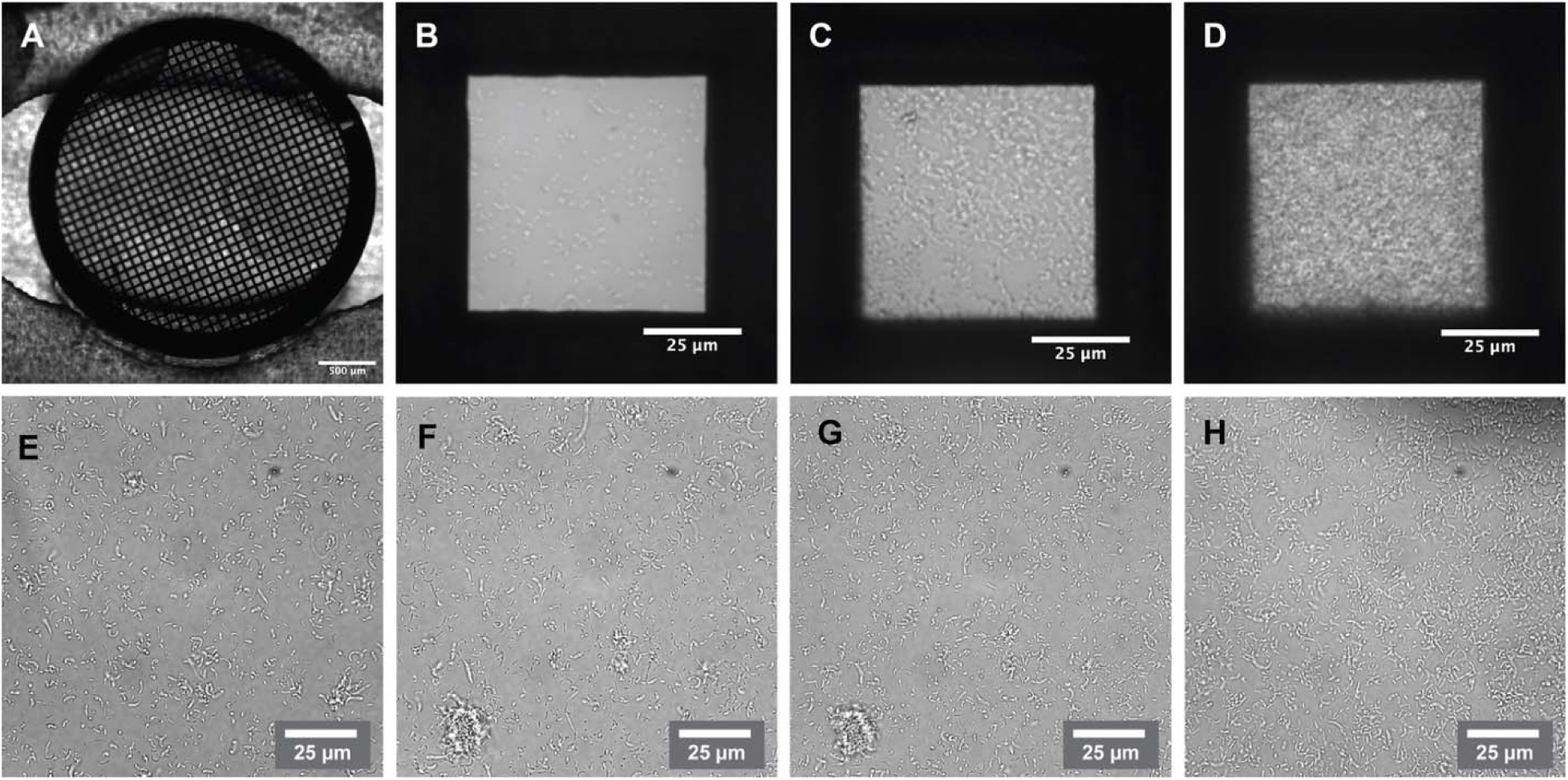
Fluid flow experiment with Caulobacter crescentus CB15 cells. *Caulobacter crescentus* CB15 cells were inoculated into the FLUID-CELL and incubated under static conditions for 2 hours to allow for grid attachment. Next, the media was flushed through at a rate of 200 µL/min, and brightfield (BF) images were collected every 24 hours over a 48-hour period. Time 0 represents the point at which the flow was initiated. (A) View of an EM grid within the receiving well. (B) 0 hours of flow, 40x BF image of a center square. (C) 24 hours of flow, the same grid square. (D) 48 hours of flow, the same grid square. (E-H) An equivalent experiment conducted with an Ibidi *µ-Slide VI*^0.5^ flow cell, cells were inoculated and allowed to incubate in static conditions for 2 hours, then media flow was started at a rate of 200 µL/min, and BF images were collected at regular time points of a central section of the flow chamber. (E) A BF image of an area of several starting biofilms, 4 hours after flow began. (F) A BF image of the exact location, with a notable biofilm colony developing, 10 hours after the flow started. (G) A BF image of the same area, with the same large biofilm colony, and other various colonies growing, 16 hours after flow started. (H) A BF image of the same area shows that the extensive biofilm had grown to a height at which the fluid flow swept it out of the frame; however, other, smaller colonies continued to develop 24 hours after the flow began. Scale bars of 500 μm (A), 25 μm (B-H).

**Figure 6.**
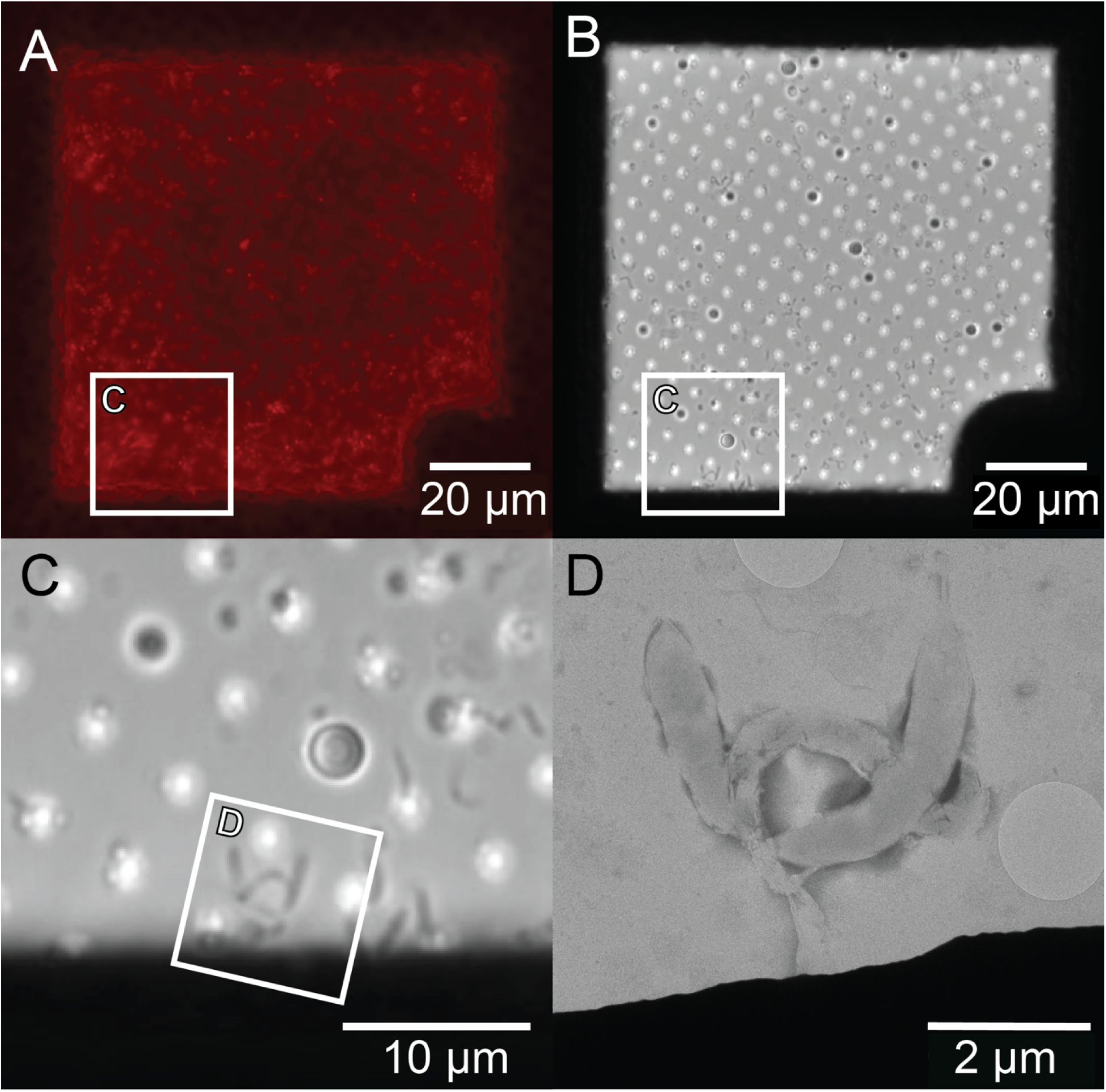
Light microscopy and electron microscopy images of *Caulobacter crescentus* cells at select time points during a flow experiment. (A) A fluorescence image of C. crescentus cells attached to an EM grid surface (whole square view) after 16 hours of fluid flow. (B) A brightfield image of the same square (A) after 16 hours of fluid flow. (C) An enlarged view of B, where a distinct ‘W’ shape of an initial *C. crescentus* cell community is visible. (D) A correlated negative stain EM image of this same *C. crescentus* cell community ‘W’. Scale bars of 20 μm (A-B), 10 μm (C), and 2 μm (D).

Our experiments highlight the relationship between flow dynamics and biological response in microfluidic systems. The use of fluorescent nanobeads to map flow profiles provided a quantitative foundation for interpreting the subsequent biofilm experiments.

By ensuring that the flow environment was well-characterized, we were able to attribute variations in bacterial behavior more confidently to the imposed hydrodynamic conditions rather than experimental artifacts. Combining precise design and engineering analysis with biological observation allowed us to create a robust methodological framework. Our approach facilitated the optimization of the microfluidic design through multiple iterations, including adjustments to channel geometry and control of flow rates. The study shows that our novel correlative microfluidic system can be readily adapted, setting the stage for advanced correlative studies that bridge the gap between cellular function and nanometer-scale architecture. This dual-modality platform offers a significant advancement for studies in structural biology and cellular dynamics.

## Summary

Manipulating EM grids, culturing cells on EM grids, and embedding physiologically relevant biological samples in vitreous ice remain technically challenging. Pipetting a culture of bacteria onto a grid and plunge-freezing them allows for the visualization of individual cells at high magnification and resolution; however, it does not support the observation of the interactions that lead to the development of a bacterial biofilm. Yet, these developmental contacts and interactions can be visualized, at lower resolution, in real-time using light microscopy combined with microfluidic systems. The design and production of synergistic technologies to support live-cell imaging, vitrification, and cryo-EM-level imaging will enhance our capacity to define in situ and ex situ architectures that regulate the function of multicellular communities.

To support broader adoption of microfluidic devices and 3D printing within cryo-CLEM workflows, we introduce FLUID-CELL, a Flow-enabled Light and Ultrastructural Imaging Device for Correlative Electron and Light Localization, as a tool for directed fluorescence light microscopy imaging of cells and cellular communities before vitrification and cryo-EM imaging. We demonstrate that 3D design software and printing equipment can support the rapid prototyping and fabrication of an EM grid-compatible microfluidic device (Figs. 1 and 2). The FLUID-CELL is designed to flexibly and securely enclose EM grids within biocompatible materials, supporting non-turbulent fluid flow across the EM grids during physiological live-cell fluorescence light microscopy experiments. Iterative design steps were employed to produce the central system, external securing bars, and internal removable gasket, rendering the device functional and leakproof for several rounds of time-intensive flow and imaging experiments of nanobeads (Fig. 4) and a model bacterium, *Caulobacter crescentus* (Figs. 5 and 6). New materials and improvements in design and fabrication technologies may permit the fabrication of additional components and other reliable flow cells for cryo-CLEM.

## Materials and Methods

### FLUID-CELL 3D Printing, Post-Processing, and Assembly

A Formlabs Form 3+ SLA 3D printer and its Preform software were used to produce the four FLUID-CELL flow cell components from three stereolithographic resins (Figs. 1 and 2). The two main body parts were printed using Clear V4 resin (25 µm layer height). The sealing gaskets were printed with Silicone 40A (100 µm layer height). The securing bars were printed with Durable V2 (50 µm layer height). The print time was approximately 2 hours for the gaskets and bars, and 10 hours for the main FLUID-CELL flow cell components.

The printed components underwent post-processing before use. The Clear V4 parts were washed with 100% isopropyl alcohol (IPA) for 30 minutes and then cured at 60 °C for 20 minutes in a Form Cure UV oven (Formlabs). The chamber bottom was machined to support the placement of an 18 mm, 1.5X, circular glass coverslip, which was secured to the frame with a double-sided adhesive and a UV-curing resin to seal the chamber edges. Silicone 40A parts were washed in 80% IPA and 20% n-butyl acetate for 60 minutes. After washing, the parts were air dried, and excess resin was removed. Once dry, the parts were submerged in water and then cured for 60 minutes at 60 °C. Durable V2 components were washed with 100% IPA for 45 minutes and then cured for 45 minutes at 60 °C. Leak testing was conducted by introducing 60 mL of liquid (ddH_2_O or PYE media) and monitoring for leakage over a 10-minute period. FLUID-CELL devices have been cleaned, re-sterilized, and reused without any adverse effects on cell culture or fluid flow. This ability is limited to ∼5 runs. After this point, continuous exposure to fluid and external pressure gradually warps the microstructure of the additive manufacturing process, resulting in larger body deformation. Once this warping reached a point where the contact between the gasket and the flow cell, or the warped geometry, breaks the glass coverslip, the FLUID-CELLs were retired from experimental use.

### Experimental Setup and FLUID-CELL Usage

To assess fluid flow dynamics and validate the functionality of the FLUID-CELL, an experimental setup was initially implemented using fluorescent beads and subsequently with *Caulobacter crescentus* cultures. Sterile tubing was assembled using Nalgene Pharma-Grade Platinum-Cured Silicone Tubing, 1/16 inch I.D (Thermo Scientific), Luer-Lok adaptors (Fisher), and Y-connectors (Fisher), as outlined (Fig. 3). The assembled tubing and media were sterilized via autoclaving (liquid cycle, 25 minutes). EM grids were glow-discharged and rinsed with ethanol before being placed and sealed into the flow cell chambers. The entire experimental system consisted of prepared inlet and outlet tubing, a programmable Gilson MINIPULS 3 peristaltic pump, a flow cell with EM grid, sterilized liquids, and bacterial culture media.

### Fluid Flow Testing

The flow characteristics of the FLUID-CELL chamber were determined by flowing sterile water containing 10^4^/mL of 1.0 μm FluoSpheres Polystyrene Microspheres (Fig. 4), with maximum excitation and emission at 578 and 607 nm, respectively (F13083, ThermoFisher), through the system while imaging using a Leica DMi8 widefield light microscope (Leica Microsystems Inc., Deerfield, IL) using a 4x lens with a numerical aperture of 0.13 connected to a Hamamatsu ORCA-Flash 4.0 camera (Hamamatsu Photonics). Movies were collected at 33.3 frames per second.

Fluorescence images were collected using the Texas Red (TXR) filter (Exc: 540-560, Ems: 630-675). This experiment was performed with multiple FLUID-CELLs to confirm uniform flow between devices. Images and movies were recorded using Leica LASX software. The movies were analyzed using ImageJ with the TrackMate plug-in^29^. Within TrackMate, the Hessian Detector^30^ was used to locate beads within each movie frame. This detector calculates a Hessian matrix for each frame, uses the determinant to identify bright concentrations, and assigns a quality score to each point based on user-defined specifications. The program was instructed to locate 8 µm diameter beads with a normalized quality score of 0.2 or higher on a scale of 0.0 to 1.0. Once this data set was collected, an additional manual thresholding was performed to ensure that detection was calibrated correctly. With this dataset of points accumulated at each movie frame, an advanced Kalman tracker was used to link particles between frames and utilize the camera’s time step to determine the velocities and streamlines of the fluid^31^. The Kalman tracker was designed for tracing particles with steady velocities; it used the velocity of a path to predict each particle’s displacement between frames and searches from this point to find the particle in the following frame. This tracer was specified to search 40 µm around each predicted displacement to continue tracing each particle. After this, further automatic thresholding was conducted to eliminate tracks that jumped between particles by either removing tracks with fewer than 5 frames or nanobeam paths shorter than 10 µm.

### Bacterial Cell Culture

*Caulobacter crescentus* strain DH1210 (*pleC*::venus, *divJ*::mkate) was a gift from Prof. David Hershey (University of Wisconsin-Madison). Bacteria were grown in peptone-yeast extract (PYE; 0.2% peptone, 0.1% yeast extract, 1 mM MgSO_4_, and 0.5 mM CaCl_2_), a medium modified from that of Schmidt and Stanier^32^.

### Bacterial Cell Culture and Flow Initiation

EM grids were glow-discharged before placement in the FLUID-CELL flow cell chamber. The assembled FLUID-CELLs with the grid were sequentially flushed with 10 mL of ethanol, sterile deionized (DI) water, and sterile PYE medium. *C. crescentus* cultures were grown in liquid PYE media ^33^ and incubated overnight. The overnight *C. crescentus* cultures were diluted to an optical density at 600 nm (OD_600_) of 0.3. The grid (and system) was inoculated with this culture and allowed to incubate under static conditions for 18 hours (∼6 growth cycles). PYE flow was introduced into the system at a rate of 200 µL/min for a specified period that depended on the experimental parameters. The grids were monitored and imaged over time using light microscopy (Fig. 5) to collect data on cell mobility and biofilm development. Once bacterial growth had reached a suitable level, the grid was removed from the chamber for downstream processing. Samples were processed and preserved by either heavy metal staining (0.5% phosphotungstic acid (PTA)) or plunge freezing using a Leica EMGP2 plunge freezer (Leica Microsystems). The grids were then imaged by TEM, and the LM and EM imaging data correlated (Fig. 6).

### Light and Fluorescence Microscopy Imaging

Light microscopy imaging was conducted using a Leica DMi8 wide-field light microscope (Leica Microsystems). Using a 63x oil immersion lens with a numerical aperture of 1.4, fluorescence images were collected with two different filter sets: FITC and TXR, both with an exposure time of 6 seconds. Images were collected with a Hamamatsu ORCA-Flash 4.0 camera (Hamamatsu Photonics) and Leica’s LASX (Leica Microsystems) imaging software. Image Z-stacks, montages, and merged maps were all collected using LASX imaging tools.

### Negative Stain EM Grid Preparation, EM Imaging and Data Processing

Grids for negative stain EM imaging were processed as follows. The grids were gently removed from the flow cell chamber, blotted, and 4 µl of 0.5% phosphotungstic acid (PTA) was applied. The grids were incubated for 45 seconds, then side-blotted with #1 Whatman filter paper (Cytiva, MA, USA) between each wash and after the final incubation with stain. Grids were washed twice with 4 ul of distilled water with blotting between washes. The grids were allowed to air dry for approximately five minutes and were stored in a desiccator until imaging. Imaging of the negative-stain grids was performed on a Talos L120C 120 kV transmission electron microscope (Thermo Fisher Scientific, Hillsboro, OR, USA) equipped with a CETA CMOS camera (Thermo Fisher Scientific, Hillsboro, OR, USA). Grid screening and data acquisition were done using standard low-dose techniques in SerialEM 3.8^34^ software. Images were collected at nominal magnifications of 45 kx (3.2 Å/pixel) and 73 kx (2.0 Å/pixel) using a range of defocus values from -0.5 µm to -2.0 µm in 0.25 µm increments.

## Acknowledgements.

We are grateful for the valuable discussions and input provided during the design phases by Dr. Daniel Parrell, Dr. Joshua Pierson, Dr. Jie Yang, Dr. Matty Gaines, Mr. Joe Baumgardt, Ms. Heather Fisher, Ms. Hayley Hirsch, Ms. Kaylee Rajek, and Ms. Ava Berdelman. This work was supported in part by the University of Wisconsin, Madison, the Department of Biochemistry at the University of Wisconsin, Madison, the Morgridge Institute for Research, and public health service grants RF1 NS110436 and U24 GM139168 to E.R.W. from the NIH. This work was supported in part by the U.S. Department of Energy, Office of Science, Office of Biological and Environmental Research under Award Numbers DE-SC0018409. We are grateful for the use of facilities and instrumentation at the Cryo-EM Research Center and the Midwest Center for Cryo-ET in the Department of Biochemistry at the University of Wisconsin, Madison. We are grateful for the computational resources supplied through the SBGrid-supported ^35^.

